# *What* and *where* in electromagnetic brain imaging

**DOI:** 10.1101/2024.04.17.589904

**Authors:** Alain de Cheveigné

## Abstract

To understand the brain, we may record electromagnetic signals and use them to infer the dynamics of brain activity (the “what”), and the location or distribution of its sources within the brain (the “where”). In this paper, these two aspects emerge from the experimental data via distinct analysis pathways. In a first step, recordings of brain activity undergo linear data-driven analysis that results in an analysis matrix, the first column of which isolates a source of interest. In a second step, the *remaining* columns of the analysis matrix are used to estimate the position of the source, with the help of a source model informed by the anatomical scans and sensor geometry. Specifically, each column is interpreted as a *null filter* for the source of interest (a spatial filter that suppresses that source). Each such null filter admits a *zero set*, or subset of the source parameter space for which the filter output is zero. Since the source of interest is necessarily within the zero sets of all of its null filters, its location can be found by taking the *intersection* of these zero sets. There is no theoretical limit to location accuracy, but practical limits may arise from noise in the data, imperfect calibration, or an incomplete or inaccurate source model. The method is illustrated with simulated and real data.

## Introduction

In Plato’s allegory of the cave, prisoners inferred the world indirectly from shadows shifting on the wall. According to Helmholtz’s concept of *unconscious inferenc*e, a similar process allows us to apprehend the world from our senses (Alhacen 1030; Helmholtz 1867; Hatfield 2002; Imbert 2020; Kawato 1993; Friston 2018). Sensory information is sparse and often incomplete (as exemplified by the shadows), and thus perception depends critically on an *internal model* of the world, constrained by the fragmentary sensory patterns. Interestingly, a similar situation arises when we try to estimate the spatial distribution of brain activity from electromagnetic (EM) data, such as electroencephalography (EEG) or magnetoencephalography (MEG). The data are too sparse to fully constrain the inverse problem, and thus estimates of source position or distribution must rely heavily on models and priors, often informed by anatomical knowledge based on MRI scans co-registered with the sensor geometry (Baillet 2001, 2010). Perception and brain imaging both rely critically on inference.

Shadows, in Plato’s metaphor, lack color and texture information, and thus they offer feeble clues as to *what* the outside objects are, but their sharp boundaries offer strong clues as to *where* the objects are located relative to the source(s) of light. Curiously, a what/where distinction is also observed between perceptual processing pathways in the brain (Schneider 1969), suggesting that there might be some functional advantage in distinct processing for the two kinds of information. Here, I draw loosely on Plato’s metaphor to motivate a new approach to the localization of sources of brain activity.

In electromagnetic (EM) brain imaging, multi-channel measurements of brain activity are used together with anatomically-informed source and forward models to infer the spatial distribution of brain sources. The source model constrains the nature and position of potential source(s), and the forward model relates activity at each location to observations at the sensor or electrode array (these models are sometimes referred to jointly as a “source model,” here they are referred to as distinct). Source and forward models are informed by anatomical measurements (e.g. from magnetic resonance imaging, MRI), knowledge about tissue conductance, sensor/electrode geometry, and other priors (e.g. regions of interest, ROI). Sophisticated software tools are available to implement such models (Oostenveld 2011; Tadel 2011; Gramfort et al 2014). Three main approaches to source localization and imaging are: *dipole fitting, beamforming*, and *minimum current norm estimation* (MNE), and here I suggest a fourth, *zero-set imaging* (ZSI).

## Methods

### Initial linear analysis

The approach involves two distinct pathways, that both stem from an initial linear analysis of the EM data. Typical analysis methods include *principal component analysis* (PCA), *independent component analysis* (ICA), *canonical correlation analysis* (CCA), *correlated components analysis* (CorrCA) (Parra et al 2019), *joint decorrelation* (JD) and other methods based on generalized eigenvalue decomposition (see de Cheveigné and Parra 2014 for a review). The objective is to find one or more *spatial filters* (columns of the analysis matrix) that isolate or enhance the time course of activity of one or more sources of interest. The methods differ according to the criterion used: variance for PCA, an empirical measure of statistical independence for ICA, correlation with another data set for CCA, contrast between two conditions for JD, and so on.

The rationale for applying linear analysis is that the EM data are themselves the result of a mixing process that is assumed to be instantaneous and linear:

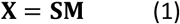

where matrices **X** and **S** represent the time series of observations and source activity respectively (**S** includes both brain and artifact or noise sources). Linear analysis involves applying an analysis matrix to the observations:

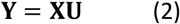

where **Y** is a matrix of time series (“components”), some of which hopefully isolate sources of brain activity. **S** and **M** are unknown, **X** is observed, and **U** is estimated by the linear analysis method.

Neural sources number in billions whereas EM channels number in tens or hundreds so, for *I* sources and *J* observation channels, *I* ≫ *J*. A useful viewpoint is that the time series of source activity span a *vector space* of dimension *I* that includes all their linear combinations. The observations *x*_*j*_(*t*) then belong to a *subspace* of dimension *J* of the source signal space. The mixing process projects brain sources onto that subspace but, since *I* > *J*, different sources may map to the same observation, reflecting the indeterminacy of the inverse problem.

Each column **u**_*k*_ of the analysis matrix constitutes a *spatial filter* (set of *J* coefficients) that, when applied to the data matrix, produces a component time series *y*_*k*_(*t*). If the analysis is successful, a column **u**_*k*_ might isolate a source of interest. By “isolate” I mean that the signal-to-noise ratio (SNR) of that source is maximized within *y*_*k*_(*t*) and moreover, importantly, *the remaining components are not correlated with that source*. The ZSI method depends critically on this property. Rather than one, the analysis might isolate a set of brain sources within a subset of components (e.g. *k* ≤ *n*), the assumption then is that the remaining components (*n* > *k*) are uncorrelated with those sources.

From this point, the analysis proceeds along two independent pathways (Fig. 1). The first applies columns **u**_*k*≤*n*_ of the analysis matrix to the EM observation matrix to estimate the time course(s) of brain activity. The other combines the remaining columns **u**_*k*>*n*_ with source and forward models to estimate the location or distribution of the source(s). This second pathway depends on the EM observations only indirectly, via the matrix **U** . Conversely, the first pathway does not involve anatomical knowledge. In this sense, we have distinct “what” and “where” pathways (Fig. 1).

**Figure 1.**
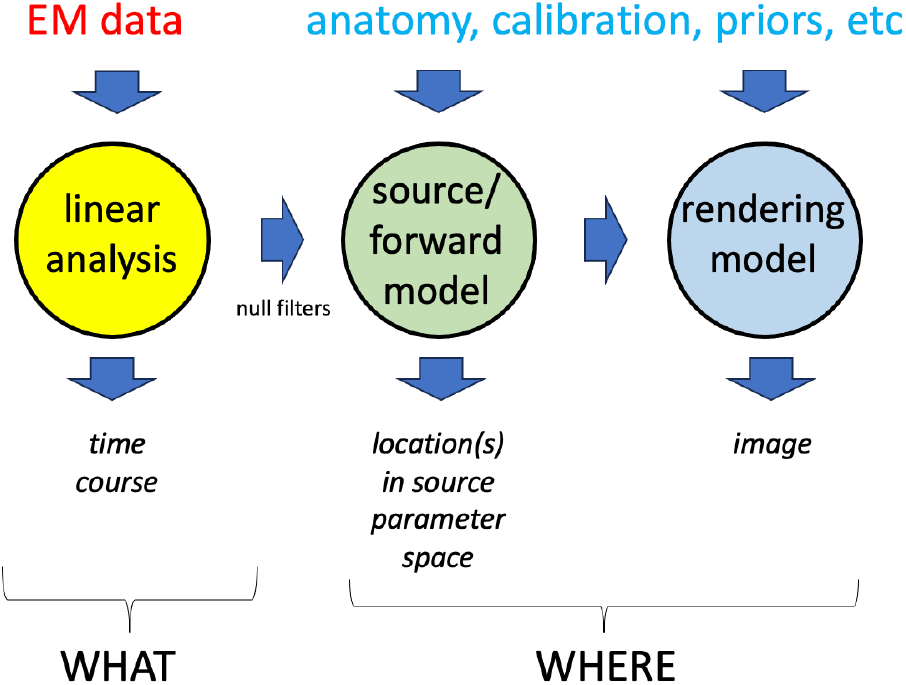
Schematic view of EM data analysis. The time course of activity of interest is derived from linear analysis of the EM data, without the help of spatial information (anatomy, etc.). The location of the source (in parameter space) is derived from source and forward models informed by spatial information, associated with null filters derived from the linear analysis. The location within source parameter space may be translated into a visual form with a rendering model, also informed by anatomy and priors.

### Source location in parameter space

The *source model* constrains the nature and location of the source. It can be understood as defining a *search space* within which to find a parameter value (“location”) that best fits the EM observations. Parameters might include 3D position, but also orientation and non-spatial features such as the nature of the source (e.g. dipole vs multiple dipoles or patch), and thus the parameter space may be distinct from physical space. For clarity I will use the term “location” to refer to a value within this parameter space, and “position” and “orientation” to refer to values in physical space.

The source model is typically informed by anatomical information (for example from an MRI scan), but it might also include additional prior assumptions (for example that the dipole orientation is orthogonal to the cortical surface, or it is positioned within a restricted region of interest). Complementary with the source model is a *forward model* that specifies the gain between every source location and each of the *J* sensors. The forward model is typically informed by both anatomical information and the geometry of the sensors, co-registered into shared coordinates. The forward model can be summarized as a forward gain matrix **M** of dimensions *I* × *J* that relates source signals *s*_*i*_(*t*) to the observations *x*_*j*_(*t*).

A spatial filter **u** is a set of *J* coefficients (column vector) that, applied to the observation matrix yields a component *y*_*u*_(*t*) = **Xu**. For a single model source *s*_*i*_(*t*), the output of the filter is

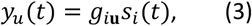

where the factor *g*_*i***u**_ is the product of the *i*-th row of **M** by the spatial filter. A filter for which *g*_*i***u**_ = 0 is called a *null filter* for a source located at *i*. A linear combination of null filters for a source is a null filter for that source, thus null filters for a source form a *vector space* which, since there are *J* coefficients and one constraint, is of dimension *J-1*. Bottom line: for any source *i*, there exist numerous spatial filters {**u**}_*i*_ that are null filters for that source.

Conversely, for a given filter **u**, there may exist a set of source parameters {*i*}_**u**_ for which **u** is a null filter. I call this set (which may be empty) the *zero set* of the filter **u**. The zero set of a null filter for a source of parameter *i* obviously includes *i*, and if several such filters are available, their intersection includes *i*. This is the principle behind zero set imaging: given (a) *a set of null filters for a particular source* and (b) *an accurate source/forward model*, the location of that source is obtained as the intersection of their zero sets.

At this point, the reader might feel confused by these various “spaces” (*signal space, parameter space, physical space*) and their significance. To ease visualization, I will introduce a “toy world” with simplified geometry and propagation properties, designed to capture the essence of the full problem while allowing for intuitive understanding.

### A “toy world

Sources and sensors are within a plane (the former within a disc, the latter on its periphery), and source-to-sensor gain varies as the inverse of the square of their distance, *m*_*ij*_ = 1/*d*_*ij*_^2^ (Fig. 2 A). The planar geometry of the source space is homeomorphic to the cortical surface, and the gain dependency roughly mimics Maxwell’s equations, while real-world complexities such as three-dimensional location and orientation, anatomical structure, and inhomogeneous or anisotropic conduction, etc. are conveniently ignored.

**Fig. 2.**
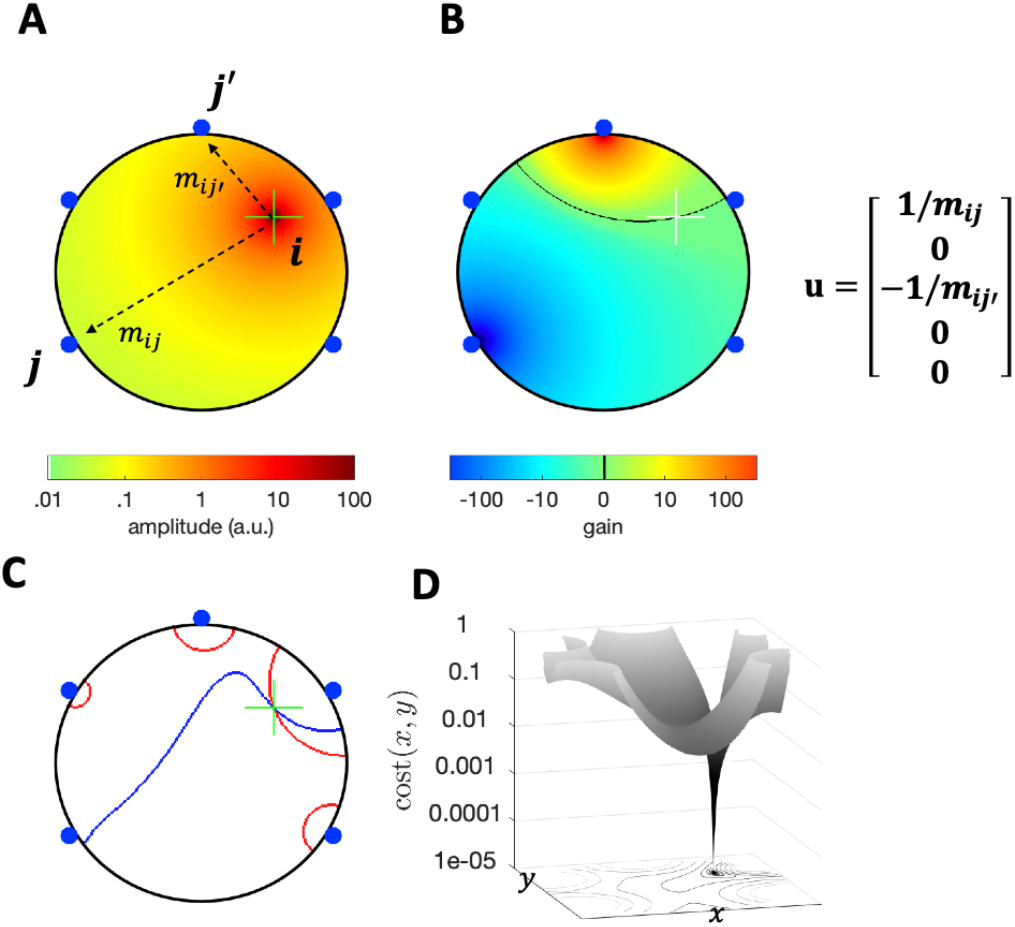
“Toy world” consisting of an array of sensors (blue dots) placed at the periphery of a disc. A: A source within the disc (green cross) radiates within the plane with an amplitude that decreases with distance as 1/*d*^2^. B: Gain map for a linear combination of two sensors with weights chosen to cancel the source at the green cross (*null filter*). The weight vector (spatial filter) is shown on the right with coefficients ordered clockwise from the lower right. The black line is the *zero set* of this filter. C: Zero sets of two other null filters for this same source. The zero sets intersect at the source. D: Cost function of position within the source space (Eq. 5) for four null filters. The global minimum (zero) also indicates the source position.

A source (green cross) radiates within the disc as shown as a color scale (Fig. 2 A), with a different gain at each sensor. A null filter **u** for that source has a gain pattern (function of source position *i*)

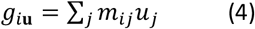

that is zero for this source (Fig. 2B). This particular null filter was obtained by setting coefficients *u*_*j*_ = 1/*m*_*ij*_ and *u*_*j′*_ = −1/*m*_*ij′*_ and all others to zero, but there exist many other null filters for that same source; the zero sets of two of them are shown in Fig. 2C. The zero set of each filter is extended in source parameter space, but they all intersect at the position of the source.

A convenient way to find the intersection is to calculate a “cost function” as

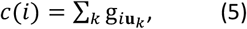

where **u**_*k*_ are null filters, and search for the value of *i* for which it is zero. Figure 2D shows a 3D plot of the value of this cost for four null filters. A deep dip is visible at the location of the source.

#### Is the zero unique?

If there are several zeros within the source space the location estimate is ambiguous. Might this occur? This question is difficult to answer in full generality, but some insights can be gained within this toy world. The zero set for filter **u** consists of positions *i* = (*x, y*) such that

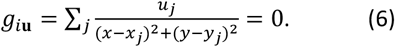

Multiplying each term by the denominators of the other terms (to give all terms the same denominator) and summing, we have:

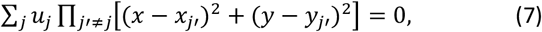

which can be expanded into a multivariate polynomial equation in *x, y* of order 2(*J* − 1). We have a similar equation for each null filter, together they form a set of simultaneous equations. Since there are two unknowns (*x, y*), we need at least two such equations, and thus at least two null filters (consistent with the geometric argument that requires intersecting zero sets).

Bezout’s theorem tells us that there may be up to [2(*J* − 1)] ^2^ solutions to this set of equations (although some might be complex and thus not admissible as coordinates). This is both good news, and bad. The bad news is that the solution is not necessarily unique, so there might indeed be situations where the location estimate is ambiguous. The good news is that their number is finite so that, in the hypothetical case where several solutions are found, we can at least enumerate them. With the geometry of Fig. 2A, the zero appears generally to be unique.

Summarizing, the steps required to locate a source within the toy world are:

1. find two or more null filters for that source,
2. given these null filters, and the source/forward model, calculate a cost function using Eqs. 4 and 5
3. search the parameter space of the source model for a zero of this cost function.

In practice, we are unlikely to find a true zero due to various sources of error, so we must search instead for a *minimum*. The cost function of Eq. 5 tends to vary widely over the search range (it approaches infinity near each sensor), which complicates the search for a minimum. This problem is alleviated by replacing Eq. 5 by

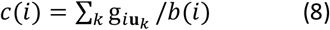

where the normalization term is calculated as,

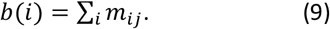

This has the effect of “flattening” the cost function over most of the search range so that the minimum is easier to locate.

In this toy world, the parameter space of a single point source maps to physical space (2D disc). However, it can be extended for example to a pair of point sources, or a patch, etc., in which case the parameter space no longer maps simply to physical space. If the parameter space is high-dimensional, attention may need to be devoted to *computational cost*.

### Rendering model

In our toy world, a location in source parameter space is readily visualizable, but the same might not be true for a more realistic model, such as a 3D dipole model with arbitrary orientation. To create pretty pictures (and earn the label “imaging”), we need to add a *rendering model* that translates a location in parameter space into a visible form. This model may include priors or assumptions, such as smoothness of the distribution of activity over cortical tissue, or expected anatomical boundaries, that enhance the *realism* of the image. However, it is important to realize that such “realism” comes at the expense of the status of the image as a prior-free observation.

### Implementation

Simulations and data analysis were performed in Matlab, using scripts available at https://github.com/adc2/ZSI. EM data analysis relied on the NoiseTools toolbox (https://github.com/adc2/NoiseTools), source/forward models were calculated using BrainStorm (https://neuroimage.usc.edu/brainstorm/), and examples are based on publicly-available data from the BrainStorm tutorials. Orientation estimation relied on code from https://www.mathworks.com/matlabcentral/fileexchange/37004-suite-of-functions-to-perform-uniform-sampling-of-a-sphere.

For the “toy world” model and the cortical surface model, the 2D parameter space was scanned exhaustively. For the cortical volume model (with unconstrained dipole orientation), the volume was scanned exhaustively (typically with 2mm cube resolution) and, for each 3D voxel, the optimal orientation was estimated as follows. A uniformly-distributed set of 100 orientations was defined (Semechko 2025), and the cost function estimated for each orientation. The orientation with smallest cost was chosen and four-point weighted interpolation was applied to it and three closest neighbors to determine a new, optimal orientation for which the cost function was estimated. The estimated cost is assigned to the 3D voxel (together with the orientation), allowing for visualization (as in Fig. 4B-D).

## Results

### Simulations

#### Toy world

A “source” waveform was synthesized as a column vector of 1000 random numbers, and multiplied by a gain vector corresponding to the position indicated by the green cross in Fig. 1A, to obtain an “observation matrix” of size 1000 ×5. PCA was applied, producing an analysis matrix of size 5 × 5, the last four columns of which were used as null filters.

Combining those filters with the source/forward model according to Eq. 8 yielded the cost function plotted in Fig. 2D.

This simulation illustrates the process by which null filters are “distilled” from the observed signals using a data-driven linear analysis method, and combined with a source/forward model to obtain the location of the source.

#### MEG source model

A source/forward model was prepared from a BrainStorm tutorial dataset for 274-channel MEG. The anatomically-informed source model allowed for a single dipole in any of 15002 positions on the cortical surface, with orientation orthogonal to that surface. The forward model was embodied by a 15002 × 274 gain matrix. A “source” was chosen with the location illustrated by a circle in Fig. 3A, and an activity waveform for that source was synthesized as a column vector of 10000 random numbers. This was multiplied by the appropriate column of the gain matrix to obtain an “observation” matrix of size 10000 × 274. PCA was applied, producing an analysis matrix of size 274 × 274, and the last 273 columns of which were used as null filters. Combining those filters with the source/forward model yielded the cost function plotted in Fig. 3B. This function shows a deep minimum at a location that matches the ground truth (compare panels A and B). This simulation extends the previous one to a more realistic configuration. In this noise-free, fully accurate simulation, the location is obtained with a spatial resolution limited only by the grid resolution of the source/forward model. Variants of these and other simulations are described in the Supplementary Information (SI).

**Fig. 3.**
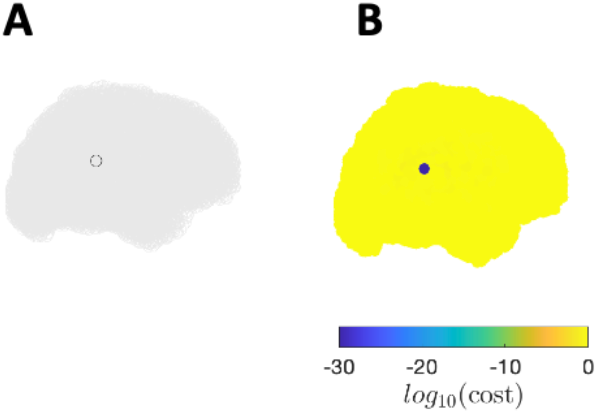
Simulation with cortical surface model. The location of the simulated source (circle in A) is recovered perfectly (blue in B).

**Fig. 4.**
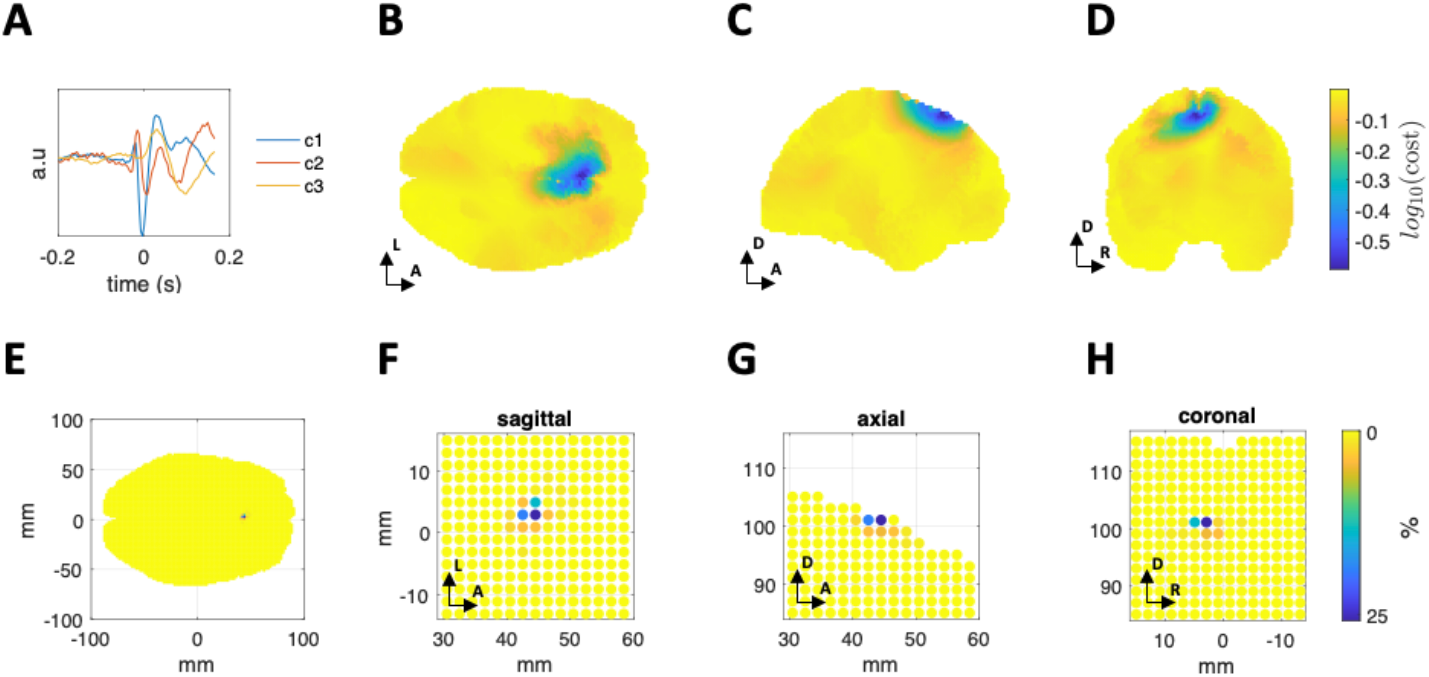
Localization of interictal spikes in EEG. A: Waveforms of three components of a JD analysis time locked to interictal spikes. B-D: Cost function associated with the first (earliest) component, based on a volume source model on a 2mm grid, projected on sagittal, axial, and coronal planes (minimum-take-all). E-H: empirical probability distribution of the position of the minimum cost, based on bootstrap resampling (n=1000).

### Real data

### Interictal spikes in EEG

The method was applied to 29-channel EEG data recorded from an epileptic patient between seizures (Brainstorm tutorial). The data, which included 58 expert-labeled interictal spikes, were first analyzed with the JD method (de Cheveigné and Parra 2014) to isolate activity time locked to markers aligned with the spikes. Figure 4A shows the time-averaged waveforms of the first three JD components which collectively span the temporal activity associated with the spike (their multiplicity reflects the presence of multiple sources with distinct spatiotemporal signatures). Singing out the first, earliest JD component (blue in Fig. 4A), and interpreting it as a single, localized source, the remaining 28 columns of the JD analysis constitute null filters for that source.

A source model (dipole with unconstrained orientation on a 2mm^3^ grid within the brain volume) was defined on the basis of anatomical information for the patient, and a forward model (source position-to-electrode gain matrix) was calculated with the help of the Brainstorm software. These null filters were associated with the source/forward model to obtain a cost function that was calculated over the brain volume, with the orientation at each voxel chosen to minimize the function. The cost function is plotted in Fig. 4B-D in a “transparent brain” representation (each pixel represents the minimum over a stack of voxels orthogonal to the viewing plane). The cost function is smallest at a superficial position on the left margin of the longitudinal fissure, roughly consistent with results from dipole fitting, beamforming, and minimum norm estimation (SI). This is the position of a dipolar source that best fits the data.

To estimate the robustness of this estimate with respect to signal variability, the estimate was repeated while sampling among spikes according to a bootstrap procedure (n=1000). As shown in Fig. 4E-H, the distribution is tight: 95% of trials fall within 4 mm of its centroid. However, in the absence of ground truth, one cannot say if that that centroid coincides with the true source. Additional examples of applying the method to real EEG, MEG, or intracranial EEG are described in SI.

A word of caution is in order at this point. The theory assumes that the source space is *complete*, in the sense that there exists a parameter for which the model source matches the real source, at which point the cost function must approach *zero*. In the example of Fig. 4, the minimum value is not zero (see color scale in panel D), and a similar outcome was observed for other data sets (SI). The possible significance of this mismatch is addressed in the Discussion.

## Discussion

This paper presented a new approach to brain source localization and imaging, described the philosophy and principle of the method, and reported results with simulated and real data. These results constitute a summary “proof of concept”, showing that the method works and is potentially applicable to real data. More work is required to validate the method and compare it to existing methods, but this is beyond the scope of this paper. The SI explores multiple aspects, however a more exhaustive investigation is warranted, including *theoretical analysis* (e.g. of sensitivity to parameter mismatches), *simulations*, and *evaluation with real data*. The rest of this Discussion is devoted to formulating the questions to ask (rather than answering them).

When the method was applied to a real recording (Fig. 4), the cost function did not reach zero for *any* value within the parameter space (colorbar in panel D), and the same was found for other analyses of real data reported in SI. At face value, this implies that the assumptions laid out in the Methods are not strictly valid. This might result from one or more of the following: (a) the source model is incomplete, for example it assumes a single dipole but the brain source is not a single dipole, or (b) the forward model is inexact, for example due to miscalibration or simplifying assumptions (e.g. conductivity), or (c) the null filters are inexact, for example due to noise in the recordings. These limitations are to be expected, the question being how serious are they. In a nutshell: is the location derived from an approximately correct model approximately correct?

The question may be amenable in part to a *theoretical* approach, to better understand the impact of each of these potential limitations. A more expedient approach may be *simulation*, for example starting with a realistic source/forward model, using it to create a source with known characteristics, and investigating the ability of the method to recover that source in the event of a model mismatch (e.g. a cortical surface-constrained dipole with an orientation that differs from the normal to the surface, or multiple dipoles, etc.), or an inaccurate forward model (e.g. due to incorrect sensor calibration), or noise in the recordings from which the null filters are derived. Of obvious interest, also, is how the method deals with real data, with the caveat that in this case the ground truth is lacking. Several examples are described in the SI.

Of particular interest is how the method is related to, and compares with, established methods such as MNE, dipole fitting, or beamforming. A feature of the new method is the clean separation that it makes between “what” and “where” pathways. Localization depends on the EM data only via the null filters, which serve as “sufficient statistics” into which the EM observations are distilled. Another feature is the unusual reliance on *cancellation* as a means to characterize a source, which bears analogy with predictive coding (Attneave 1954; Barlow 2001; Friston 2018), and cancellation-based theories of sensory processing (Durlach 1963; de Cheveigné, 2023a) and, ultimately, with Plato’s metaphor.

This method, like others, relies on strong priors to complement the extremely sparse observations (Baillet 2001, 2010). A possible appeal is the clear modularity between observations (encapsulated in null filters), source model, forward model, and rendering model. This is in contrast to e.g. MNE for which a smoothness prior is implicit in the minimum norm constraint. Also in contrast to MNE, the pattern formed by the cost function over source space (e.g. Fig. 4B-D) is *not* interpretable as a pattern of activity. Rather, it quantifies the pattern of *fit* over the model parameter space. This is a potential downside of this new method, as it is extremely tempting to interpret e.g. Fig. 4B-D as indicating spread of activation. Also tempting is to interpret it in terms of *probability*: that link too is not established.

Granted accurate source and forward models, and accurate null filters, there is no limit to spatial resolution beyond that of the grid used to sample the source space. This is in contrast to the concept of point spread function applied to other methods (Sato 2004; Lütkenhöner 2003; Jaiswal et al 2020; Hauk et al 2022). A nice feature is that, while it clearly distinguishes activity and location pathways, the new method draws together data-driven signal-enhancement approaches (such as ICA) and localization approaches that are rarely discussed together other than in terms of, e.g., using ICA for artifact rejection. Here, enhancement and localization both stem from the initial linear analysis.

ZSI resembles *dipole fitting* in that the source model parameter space is scanned for a fit to the data. In dipole fitting, the fit is made directly to values observed at the sensors at a certain time point, and thus it is vulnerable to competing sources and noise. In ZSI, this is alleviated by the initial linear analysis that factors the data into distinct subspaces. *Beamforming* too involves a scan of the source parameter space for a solution that best enhances the source of interest (according to some criterion applicable to the beamformer output). Like ZSI, it relies on cancellation, but whereas ZSI focuses on cancelling the target, beamforming cancels the interferers, or a subset thereof. As a localization method, beamforming is subject to a “switching” behavior, best understood in a hypothetical scenario with *J* sources for *J* sensors. As the source space is scanned, the beamformer switches between solutions involving the cancellation of *J* − 1 sources, each of which is unique (de Cheveigné 2023c). The source space is thus parcellated into *J* − 1 regions, implying in that case a crude spatial resolution (see SI for relevant simulations).

In contrast to ZSI and other methods that scan the source space, MNE directly estimates an “inverse” matrix **U** using a constraint of minimum current norm to overcome indeterminacy. Applying that matrix to the observations **X** yields a “reconstructed” source activity matrix 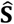 that packs both “what” and “where” information. An advantage is that the minimum norm constraint enforces a prior that leads to plausibly smooth distributions of estimated activity. However, one might be concerned that the resulting “image” reflects as much the prior as the data, which is possibly problematic if it is to be used as an “observation” for the purpose of inferring the spatial extent of neural processing circuits. In ZSI, a smoothness prior, if used, would be encapsulated in the rendering model, while anatomical priors would fit within the source model.

A weakness of the new method is the assumption of a model source space that is complete (i.e. there exists a parameter that matches the true source). Most source/forward models assume a single dipole (more or less constrained), which might not match a source that is extended (particularly on a non-planar surface) or multifocal (e.g. both hemispheres). The elementary dipole model can be extended, but this poses issues that are both practical (search cost) and theoretical (overfitting). On the bright side, the minimum value of the cost function is a reliable “canary in the coal mine” to indicate that the source model is *not* complete (assuming forward model and null filters are exact).

In neuroscience it is common to speak of distinct what/where pathways within the brain (Schneider 1969), a distinction that might reflect computational constraints (Chikkerur 2010; Cisek 2019). The method is applied here to brain data, but it might be more widely useful, e.g. in acoustics or geophysics. Based on the reciprocity theorem, sensor positions can in principle be localized from a calibrated array of sources, thus contributing to instrumentation. These and other aspects remain to be explored.

Returning to Plato’s allegory, if the shape of the shadow were scaled and turned into a mask placed *between* the light source and the object, it would perfectly obliterate that object. This is an almost perfect metaphor for a null filter. Just as multiple light sources cast multiple shadows of the same object, together informing us of its volume in space, multiple null filters constrain the location of a source within the brain. Looking at an illuminated object tells us *what* that object is, and looking at the shadows that it casts tells us accurately *where* it is, an idea that works also for the brain.

## Acknowledgment

Previous drafts benefitted from comments of Israel Nelken, Lucas Parra, Carsten Wolters, and Malcolm Slaney.

## References

al Haytham I. (1030) Book of optics (see Hatfield 2002, Imbert 2020).

Attneave, F. (1954). Some informational aspects of visual perception. Psychological Review, 61(3), 183–193. 10.1037/h0054663

Baillet, S., Mosher, J.C., and Leahy, R.M. (2001) Electromagnetic brain mapping. IEEE Signal Processing Magazine, 18(6):14–30.

Baillet, S. (2010) The Dowser in the Fields: Searching for MEG Sources, in MEG, an Introduction to Methods, edited by Hansen, P.C., Kringelbach, M.L., Salmelin, R., Oxford University Press, 83–123.

Barlow, H. (2001). The exploitation of regularities in the environment by the brain. Behavioral and Brain Sciences, 24(4), 602–607. 10.1017/S0140525X01000024

Barnes, G. R., Hillebrand, A., Fawcett, I. P., & Singh, K. D. (2004). Realistic spatial sampling for MEG beamformer images. Human Brain Mapping, 23(2), 120–127. 10.1002/hbm.20047

Chikkerur, S., Serre, T., Tan, C., & Poggio, T. (2010). What and where: A Bayesian inference theory of attention. Vision Research, 50(22), 2233–2247. 10.1016/j.visres.2010.05.013

Cisek, P. (2019). Resynthesizing behavior through phylogenetic refinement. Attention, Perception, & Psychophysics, 81(7), 2265–2287. 10.3758/s13414-019-01760-1

de Cheveigné, A., Parra, L. (2014) Joint decorrelation: a versatile tool for multichannel data analysis. NeuroImage, 98, 487–505, doi: 10.1016/j.neuroimage.2014.05.068.

de Cheveigné, A. (2023a) In-channel cancellation, a model of early auditory processing, J. Acoust. Soc. Am., 153, 3361–3372, 10.1121/10.0019752.

de Cheveigné, A. (2023b) Predictive coding in the auditory brainstem, bioRxiv, 10.1101/2023.12.31.573202.

de Cheveigné, A. (2023c). Virtual electrode or virtual scalpel? A cancellation-based approach to data analysis and brain source localization, bioRxiv, 10.1101/2023.09.19.558390.

Durlach, N. (1963). “Equalization and cancellation theory of binaural masking-level differences,” J. Acoust. Soc. Am. 35, 1206–1218.

Frauscher, B., Von Ellenrieder, N., Zelmann, R., Doležalová, I., Minotti, L., Olivier, A., Hall, J., Hoffmann, D., Nguyen, D. K., Kahane, P., Dubeau, F., & Gotman, J. (2018). Atlas of the normal intracranial electroencephalogram: Neurophysiological awake activity in different cortical areas. Brain, 141(4), 1130–1144. 10.1093/brain/awy035

Friston, K. (2018). Does predictive coding have a future? Nature Neuroscience, 21(8), 1019–1021. 10.1038/s41593-018-0200-7

Gramfort, A., Luessi, M., Larson, E., Engemann, D. A., Strohmeier, D., Brodbeck, C., Parkkonen, L., & Hämäläinen, M. S. (2014). MNE software for processing MEG and EEG data. NeuroImage, 86, 446–460. 10.1016/j.neuroimage.2013.10.027

Hatfield G. (2002) Perception as unconscious inference. In Heyer D., & Mausfeld R. (Eds) Perception and the physical world: Psychological and philosophical issues in perception (pp. 113–143). John Wiley and Sons.

Hauk, O., Stenroos, M., & Treder, M. S. (2022). Towards an objective evaluation of EEG/MEG source estimation methods –The linear approach. NeuroImage, 255, 119177. 10.1016/j.neuroimage.2022.119177.

Helmholtz H. (1867). Handbuch der Physiologischen Optik (English tranl.: 1924 JPC Southall as Treatise on Physiological Optics) Voss.

Imbert M (2020). La fin du regard éclairant. Une révolution dans les sciences de la vision au XIe siècle. Ibn al-Haythan, Vrin, Paris.

Jaiswal, A., Nenonen, J., Stenroos, M., Gramfort, A., Dalal, S. S., Westner, B. U., Litvak, V., Mosher, J. C., Schoffelen, J.-M., Witton, C., Oostenveld, R., & Parkkonen, L. (2020). Comparison of beamformer implementations for MEG source localization. NeuroImage, 216, 116797. 10.1016/j.neuroimage.2020.116797

Kawato, M., Hayakawa, H., & Inui, T. (1993). A forward-inverse optics model of reciprocal connections between visual cortical areas. Network: Computation in Neural Systems, 4(4), 415–422. 10.1088/0954-898X_4_4_001

Lütkenhöner, B. (2003). Magnetoencephalography and its Achilles’ heel. Journal of Physiology-Paris, 97(4–6), 641–658. 10.1016/j.jphysparis.2004.01.020

Oostenveld, R., Fries, P., Maris, E., Schoffelen, J.-M. (2011) FieldTrip: Open Source Software for Advanced Analysis of MEG, EEG, and Invasive Electrophysiological Data. Computational Intelligence and Neuroscience, 2011:156869.

Parra, L. C., Haufe, S., & Dmochowski, J. P. (2019). Correlated Components Analysis—Extracting Reliable Dimensions in Multivariate Data. Neurons, Behavior, Data Analysis, and Theory, 2(1). 10.51628/001c.7125

Tadel, F., Baillet, S., Mosher, J. C., Pantazis, D., & Leahy, R. M. (2011). Brainstorm: A User-Friendly Application for MEG/EEG Analysis. Computational Intelligence and Neuroscience, 2011, 1–13. 10.1155/2011/879716

Schneider, G. E. (1969). Two Visual Systems. Science, 163, 895–902.

