## Supplementary Information for "*What* and *where* in electromagnetic brain imaging"

last edited 2025 March 23

[To editor and reviewers: this material will be pruned before publication. I appreciate any suggestions as to what should be removed, or kept as essential.]

### What and where in electromagnetic brain imaging – supplementary information

#### Supplementary methods

##### Data and analysis model

Recorded data are represented as a matrix $\mathbf{X}$ of size $I \times T$, related to brain sources via a linear mixing process:

$$\mathbf{X} = \mathbf{SM} \qquad \text{(S1)}$$

where $\mathbf{M}$ is a matrix of size $I \times J$, with $I \gg J$. This equation does not include a noise term: noise sources are assumed to belong to $\mathbf{S}$. The matrix $\mathbf{X}$ is observable, $\mathbf{S}$ and $\mathbf{M}$ are not.

It is useful to reason in terms of *vector space*, with useful concepts such as *subspace*, *orthogonality*, *linear dependence*, *projection* and *dimensionality*. The columns of $\mathbf{X}$ span a subspace of the larger vector space of signals spanned by the sources**.** Different source signals $\mathbf{s}_i$ may project on to the same observation $\mathbf{x}_j$ (Fig. S1), which explains the indeterminacy of the inverse problem.

For analysis, a matrix $\mathbf{U}$ of size $J \times K$ is applied to the observed data:

$$\mathbf{Y} = \mathbf{XU} \qquad \text{(S2)}$$

where $\mathbf{U}$ is a matrix of size $J \times K$, each column of which defines a *spatial filter*, that is, a set of $J$ weights applicable to the columns of $\mathbf{X}$ to obtain one column of $\mathbf{Y}$. Each column of $\mathbf{Y}$ (time series) is referred as a *component*. Columns of $\mathbf{Y}$ belong to the subspace spanned by $\mathbf{X}$.

The set of all spatial filters applicable to $J$-channel data likewise forms a vector space of dimension $J$. Among filters, those that suppress a given source $\mathbf{s}_i$ (i.e. filters $\mathbf{u}$ such that $\mathbf{s}_i\mathbf{M}\mathbf{u} = 0$) form a subspace of dimension $J - 1$. A filter that suppresses a source is called a *null filter* for that source. Filters that also suppress a second source form a subspace of dimension $J - 2$, and so-on. Filters that suppress $J - 1$ sources form a subspace of dimension 1 (i.e. the filter is unique to a multiplicative constant).

#### Data-driven linear analysis

The analysis matrix $\mathbf{U}$ in Eq. S2 might be predetermined, but it is usually derived from the data for example using PCA, ICA, CCA, JD, or other linear analysis methods. Typically, the purpose is to isolate a source of interest to observe its time series, thanks to a particular column $k$ of the analysis matrix. Here, instead, the focus will be on the *other* columns, $k' \neq k$, that form null filters for that source. In some situations, a group of $N$ sources may be of interest, isolated by a subset of $N$ columns of $\mathbf{U}$, in which case the $K - N$ remaining columns serve as null filters.

#### Source and forward model

The *source model* specifies the kind of brain sources expected (e.g. dipole, patch, etc.), and the space of their parameters. It may also embody prior knowledge (e.g. ROI). Source positions in this space are indexed by $\tilde{\imath} = 1 \cdots \tilde{I}$. The *forward model* then is embodied by a matrix $\mathbf{G}$ of size $\tilde{I} \times J$. The row index $\tilde{\imath}$ of $\mathbf{G}$ designates a position in model source space, whereas the row index $i$ of $\mathbf{M}$ designates an actual source in the brain.

Source and forward models are informed by the anatomy of the subject, as captured by structural MRI or other structural images, and fiducial measurements that allow to co-register anatomical and sensor coordinates. In the literature, these two models are often subsumed under the single term “head model”, here they are distinguished for clarity.

#### Localization and imaging

*Localization* of a source involves finding the best fit within the parameter space of the source model. That space might map to physical space for a simple source model (e.g. single dipole), but in other cases it is more abstract (e.g. multiple dipoles or patches). Because of the abstract nature of the source space, imaging may require an additional *rendering* model, that translates a location in source parameter space to a visual image.

#### Zero set of a filter, intersection of zero sets

Given a filter $\mathbf{u}$, the source locations for which $\mathbf{Gu} = 0$ form the *zero set* of that filter. The zero set is a subset of the source parameter space (in contrast to the kernel of the transform $\mathbf{G}$) and may have a curvilinear shape (for example as illustrated in Fig. S2). If $\mathbf{u}$ is a null filter for a source, that source's location obviously belongs to the zero set of $\mathbf{u}$, and if several null filters are known, the location is within the *intersection* of their zero sets (Fig. S2). If the intersection is unique, the location is determined, otherwise it is ambiguous. Conditions in which it might not be unique are discussed below.

#### Search

Given null filters for a source, that source's location can be found by searching for the intersection of the zero sets of those filters. In practice, one may either:

1. search for the zero sets and take their intersection (as in Figs. 2 C or S2 C), or
2. search for a zero (or minimum) of a cost function such as:

$$o(\tilde{\imath}) = \sum_k (\mathbf{g}_{\tilde{\imath}} \cdot \mathbf{u}_k)^2 / b_{\tilde{\imath}} \qquad \text{S3}$$

where $\mathbf{g}_{\tilde{\imath}}$ is the vector of source to sensor gains and $b_{\tilde{\imath}}$ is a position-dependent normalization factor. The first formulation is conceptually simpler, the second is more flexible.

The normalization factor in Eq. S3 is designed to compensate for systematic variations with relative orientation of a dipole, or depth or proximity to a sensor. If the source fits the model accurately (i.e. there exists $\tilde{\imath}$ for which Eq. S3 is satisfied), that factor is unnecessary because it does not affect the position of the zero. However, if that fit is *inaccurate*, it may be necessary to rely on the position of the minimum, and the normalization factor is essential in that case. See Fig. S2 D,F for an illustration.

For the results reported in the main paper and below, the factor was calculated as $b_{\tilde{\imath}} = \sum_k (\mathbf{g}_{\tilde{\imath}} \cdot \mathbf{v}_k)^2$ where $[\mathbf{v}_k]$ consists of *all* columns of the data-driven analysis matrix $\mathbf{U}$, whereas $[\mathbf{u}_k]$ consists of all columns *except* the first $N$, where $N$ is the number of sources being localized.

#### Simulation within a "toy world"

The three-dimensional shape of the brain and sensor configuration is not easy to visualize, and thus it is not easy to grasp the physics of the problem. For that reason, I introduce a simpler two-dimensional "toy world" in which sources and sensors are parametrized with their location $(x, y)$ within a disc in a 2D plane, and source-to-sensor gain varies with their distance as $1/d^2$ (Fig. 2A). The 2D space is homologous to a cortical surface and the $1/d^2$ dependency is analogous to that which holds for EEG and MEG (Hämäläinen et al 1993; Nunez and Srinivasan 2006). Based on the toy world, it is easier to understand results found with more complex simulations and real data.

#### Simulation with a realistic MEG source/forward model

Brainstorm (Tadel et al 2011) was used to create a source/ forward model based on publicly available sample data from the Brainstorm tutorial (https://neuroimage.usc.edu/brainstorm/DatasetIntroduction). MEG data were recorded in an auditory oddball experiment with a 274-channel MEG system, the subject's brain anatomy was estimated based on an MRI scan, and the head shape was digitized to allow coregistration between MRI data and MEG sensors. Three source models were prepared using the BrainStorm software:

1. A constrained cortical surface dipole model with 15002-dipoles oriented perpendicular to the cortical surface, parametrized by their 3D coordinates and orientation. The forward model was embodied by a 15002×274 matrix of gains.
2. An unconstrained cortical surface model with 15002 dipoles with arbitrary orientations, located on the cortical surface. The forward model was embodied by a 15002×274×3 matrix of gains.

3. A volume model with 192913 dipoles with arbitrary orientations, one for each $(2mm)^3$ within the brain. The forward model was embodied by a 192913×274×3 matrix of gains.

Based on these source and forward models, simulated MEG data were produced by choosing one or more locations in source space, and multiplying a "source" time series by the appropriate rows of the gain matrix.

#### Analysis of real MEG brain data

Source and forward models were the same as for the simulation. MEG recordings from the publicly-available data were preprocessed by applying the ZapLine algorithm to remove power line interference (de Cheveigné 2020) and the Shared Component Analysis (SCA) algorithm to reduce dimensionality from 274 channels to 70 components (de Cheveigné 2021). Linear data-driven analysis was used to isolate three types of activity: an auditory response time-locked to the stimulus, a motor preparation response time-locked to the button press, and narrowband alpha activity. More precisely, the JD algorithm (de Cheveigné and Parra 2014) was applied with three different contrasts:

1. Stimulus-locked trial-averaged data contrasted with the unaveraged data.
2. Button-press-locked trial-averaged data contrasted with the unaveraged data.
3. 10-Hz narrowband-filtered data contrasted with unfiltered data.

See de Cheveigné and Parra (2014) for further background on such analysis. In each case, the analysis produced a 274 × 70 matrix (the second dimension is limited by the intial dimensionality reduction). The first few columns of this matrix capture the time course of the activity of the sources of interest, and the remaining columns form null filters from which their location can be estimated. More details are provided with the results below.

#### Analysis of real EEG brain data

Brainstorm was used to create a volume head model for the dataset used by the Brainstorm tutorial on EEG and epilepsy. 29-channel EEG were recorded from an epileptic subject, with labels indicating the occurrence time of 58 interictal spikes. These data were accompanied by anatomical measurements (MRI) from which was derived a head model with 157666 dipoles with arbitrary orientation distributed on a 2 mm grid in the cortical volume. Null filters were derived from a JD analysis designed to isolate activity time-locked to the spikes.

#### Computation

Simulations were performed in Matlab using routines from the NoiseTools toolbox (http://audition.ens.fr/adc/NoiseTools/), FieldTrip (Oostenveld 2011), Brainstorm (Tadel et al 2011), and the S^2 sampling toolbox (https://www.mathworks.com/matlabcentral/fileexchange/37004-suite-of-functions-to-perform-uniform-sampling-of-a-sphere).

For the toy world simulations, the cost function was calculated explicitly over a grid of source positions. The estimated source location was derived from the position of the minimum over this grid or, in some cases more accurately using the matlab routine fminsearch() based on the Nelder-Mead simplex algorithm.

For the simulated and real MEG analyses and the constrained-dipole model, the cost function was calculated explicitly for all dipole locations. For the unconstrained dipole models (cortical surface or volume), for each position within the 2mm grid, the orientation of the dipole was calculated for 100 orientations uniformly distributed over the sphere, and the final orientation estimate chosen by weighted interpolation between the 4 best orientations.

For the two-dipole source model (based on the constrained dipole source model), positions were pruned by first applying a one-dipole model and selecting positions with lowest cost function (e.g. ~1000). The two-dipole cost function was then calculated over the full parameter space of the two-dipole model.

### Supplementary results

#### Toy world

The purpose of the toy world simulations is to gain insights about the zero-set method. That method is conceptually quite different from others, and raises many questions that may be easier to answer within the context of the toy world than within a realistic simulation with a complex 3D structure.

The basic setup is shown in Fig. 2 A with $J$ =5 sensors. Given a spatial filter (a vector of $J$ coefficients), the filter output depends on the position of the source. The source-to-filter gain for one simple 2-coefficient filter is shown in Fig. 2 B as a color scale, that for another filter is shown in Fig. S2 A as a 3D plot with color. For the latter, the gain is strongly negative near three sensors, and strongly positive near three others, with a gradient between them. The squared gain of this same filter is shown in Fig. S2 B, plotted as a 3D plot with a gray scale. The valley indicates the zero set of this filter, visible also as a black line in Fig. S2 A).

The zero sets of five such filters are shown in Fig. S2 C (see also Fig. 2 C). They appear curvilinear, sometimes split into several pieces. All zero sets pass by the same point, and a source at that position would thus be suppressed by all filters, reflected also by the fact that the sum of squared gain maps of all filters is zero at that point and non-zero elsewhere (Fig. S2 D, see also Fig. 2 D). Normalization (Eq. 3, Supplementary methods) produces a pattern that is smoother (Fig. 2 D), without peaks near the sensors. Such normalization is crucial for MEG (see below) to avoid the effects of source depth and orientation that otherwise strongly affect the gain maps.

**How are null filters found?** If the source location is unknown, but a sensor signal recording is available, null filters may be derived using a data-driven linear analysis method such as PCA, ICA, CCA, JD or others. The principle is as follows. *If* the data-driven method succeeds producing an analysis matrix $\mathbf{U}$ such that the desired source is isolated by one column $\mathbf{u}_k$ of the analysis matrix, *and* if the columns are mutually orthogonal, as is often the case, *then* the other columns $\mathbf{u}_{k'\neq k}$ are all null filters for that source. It is not necessary for the target source to be perfectly isolated within a component $k$ (for example activity from other sources might leak in to that component). It is only necessary that it be excluded from the other components, $k' \neq k$.

**PCA.** If the source of interest is the only one active (or its activity dominates that of the other sources), PCA will yield an analysis matrix such that the first column $\mathbf{u}_1$ captures all of the target source's activity. The other columns then are all null filters for that source. This is how Figs. 2 CD and S2 CD were obtained. However, PCA will not help if the source of interest does not dominate the variance of the data.

**ICA** produces a transform that maximizes some measure of statistical independence between columns $k$ of the transformed data. **CCA** produces a transform that maximizes correlation with some other data, for example representing the stimulus or the brain activity of a different subject experiencing the same situation (ICA and CCA are not used here).

**JD**, also known as Denoising Source Separation (DSS) or Common Spatial Patterns (CSP, see de Cheveigné and Parra 2014 and references therein), finds a transform that maximizes the variance ratio between two conditions or filtered versions of the data. The principle is illustrated in Fig. S3. Synthetic data were produced by the following steps:

1. 64-channel EEG data recorded in response to a repeated tone (Fuglsang et al 2017) were submitted to a JD analysis to maximize trial-locked activity.
2. The first component isolated by that analysis was used as the waveform of one source (the target). Four other “sources” (competitors) were obtained by applying a random $63 \times 4$ mixing matrix to the remaining components. All sources were scaled for the same variance.
3. These five sources were mixed, using a forward matrix calculated for five source positions within the disc, to obtain five sensor waveforms (synthetic data).

These synthetic data were then analyzed, again by the JD algorithm, to produce a matrix $\mathbf{U}$ such that $\mathbf{u}_1$ isolates the trial-locked activity, and columns $\mathbf{u}_{2 \ldots 5}$ were used as null filters to locate the source.

Source locations are plotted as crosses in Fig. S3 A (the blue cross is the target). The waveforms of several trials of the target are plotted in Fig. S3 B (top) as a color scale, and several trials of one of the non-target sources in Fig. S3 B (bottom). The cost function for columns $\mathbf{u}_{2 \dots 5}$ of the analysis matrix is plotted in Fig. S3 C as a gray scale. The location of the minimum (red cross) coincides with that of the source (green cross).

**What if there are more sources than sensors?** In the previous example, there were 5 sources and 5 sensors, so the target was linearly separable. Fig. S3 D illustrates a situation where $I > J$. Since the target is no longer linearly separable from competing sources, the second JD analysis cannot not isolate it perfectly (not shown), but localization of the target nonetheless succeeds (Fig. S3 D). This suggests that source *localization* might be more resilient than source *separation*.

**Is it possible to localize more sources than sensors?** The previous example involved a single target, this example investigates a situation where a larger number are to be localized (81 sources for 5 sensors). Figure S4 A shows their positions within the disc, and Fig. S4 B (top) shows their time course. Crucially, source activations are sparse, with only one source active at a time. These sources were mixed by applying an 81×5 random matrix to form a 5-column data matrix (Fig. S4 B bottom). JD was applied repeatedly to these data, each time with the objective of isolating one of the transient sources, and each time columns 2-5 of the analysis matrix were used to localize that source. The result (Fig. S4 C), albeit not perfect, shows that the method can potentially localize more sources than sensors. This suggests that it might be of use to accurately locate epileptic activity from EEG, MEG, or sEEG, or multiple neurons from invasive electrode arrays. In another simulation (not shown), the 81 sources were *spectrally* sparse (sinusoids with slowly-varying envelopes), rather than temporally, again with successful localization.

**Is the location cue always unique?** Strictly speaking, the answer is no (although it often is in realistic situations, see below). For the toy world, it was pointed out (main paper) that the cost function could have up to $[d(J-1)]^{d}$ zeros, where $J$ is the number of sensors and $d$ the number of dimensions (2 for the plane).

Figure S5 A illustrates one situation in which the location cue was not unique. A single source located within the disc was mixed with a 1×15 forward matrix into 15 sensor signals. For certain positions of the source within the plane, a “ghost” zero appears, outside the array of sensors (for a real source inside) or inside the array (for a real source outside).

Figure S5 B illustrates another such situation. Here, three sources were simulated with Gaussian noise and mixed with a 3×15 forward matrix into 15 sensor signals. PCA was applied, and columns 4-15 used as null filters. The cost function has three zeros, on the basis of which those sources can be localized as a group.

**Source-model-informed component analysis**. Interestingly, in the previous example, PCA did not separate the sources: each PC is some unknown combination of sources. However, once the sources have been *localized*, the waveform of each source can be isolated by applying null filters to the other two sources. This implies that entangled components can be disentangled on the basis of the source model (analogous to the use of a statistical independence measure for ICA).

**What if the real source does not fit the model?** In all previous examples, the source model space included a value that fit the real source. In Fig. S6, the source model assumes a single point source (as in previous examples), but the real source is active at two points. In panel A, those points coincide (equivalent to a one-point source): the cost function has a deep minimum that indicates the location. In panel B, the points are placed symmetrically (green crosses): the function shows no minimum at either location. In this situation, where the real source does not fit the model, it is *not* possible to say anything about the source location.

There may however be some tolerance if the two points are close. Panel C shows the depth of the minimum as a function of the location of the second point along a horizontal line passing through the crosses. When the points are distant, the minimum is shallow. When they are close, it is deeper (albeit less deep than for coincident points, arrow), with a position intermediate between the real points (not shown).

An incomplete source model may be *augmented* to include additional source types. In Fig. S6 D, the one-point source model with parameters $(x, y)$ is replaced by a two-point model with parameters $(x_1, y_{1,} x_2, y_2)$. In this illustration, one point of each source is constrained to the left cross, and the cost function is plotted as a function of the position of the second point. The cost is minimal when the second point of the source model coincides with the right cross, i.e. when the real source fits the two-point model. If the left-hand location were not known, both locations could be found simultaneously by search within the 4-parameter space of the two-point model.

#### Simulated MEG

Brainstorm was used to produce source and forward models for MEG based on the anatomy of a subject (example data for Brainstorm tutorial), as described in the supplementary methods. Three source models were considered: “constrained”, “unconstrained” and “volume” (supplementary methods). One or more time series of “source” activity were either synthesized from white Gaussian random noise, or assembled from real MEG data. Given source locations chosen arbitrarily, the corresponding gain matrix was applied to obtain simulated MEG data. In some simulations the signal-to-noise ratio and inter-source correlation were manipulated, as described.

Figure S7 considers the case of a single source located at the left-most position within the cortical model (dot in Fig. S7 A). The source was excited with random Gaussian noise, multiplied by the appropriate gain vector to obtain a 274-channel data matrix, PCA was applied, and the last 273 columns of the PCA transform matrix, null filters for that source, were used to calculate the cost function (Eq. S4) over source locations (Fig. S7 B). The cost function shows a very sharp and deep minimum at the location of the source. The simulation was repeated placing the source at each of the 15002 positions within the source space, and the source was correctly recovered each time. This simulation demonstrates that the localization method works within a realistic MEG recording configuration.

**What is the effect of noise?** Independent Gaussian noise was added to the simulated MEG data with equal amplitude in all channels. Figure S7 C shows the gain function of position for

SNR=0.01 (source power 1% of noise power). The depth of the minimum is reduced (note the color scale) but nonetheless clear. For SNR=0.01 or better the location estimate was correct for all source positions, for SNR=0.001 or worse it was *incorrect* for all positions.

**Does it work for more realistic data?** MEG data recorded in response to repeated auditory stimulation (Brainstorm tutorial data) were processed by the JD algorithm to enhance stimulus-locked activity. The first JD component was used as the time series of the target source, and that component was projected out from the original data to obtain a 274-channel "noise" matrix. To remove any residual time-locked structure from the noise matrix, the Fourier transform was applied to each channel, the phase of each frequency augmented by a random value (same for all channels), and the inverse transform applied. The target was multiplied by the appropriate gain vector and added to the noise with an SNR that matched the estimated SNR of the original data (SNR=0.053). These data were then submitted to a second JD analysis designed to recover this target. The last 273 columns of the transform matrix were used as null filters, leading to a cost function displayed in Fig. S7D. Again, the location of the source is clear from the position of the minimum. From this simulation, we can be confident that *if* the source of the stimulus-evoked response in the real data were a simple dipole, located on the cortical surface postulated by the source model, and normal to that surface, we would be able to localize it accurately.

**What if the source does not fit the model?** Figure S8 considers the case of a two-dipole source (dots in Fig. S8 A). The same Gaussian noise time series was used for both dipoles, and the 274-channel data matrix obtained by multiplying that time series by the gain vectors for both dipole positions and adding the result. As before, null filters were derived from the last 273 columns of the PCA matrix. The resulting cost function is shown in Fig. S8 B. Here, a minimum is observed for *one* position but not the other. Worse, if the same simulation is repeated for a random sampling of dipole pairs, the minimum coincides with *neither* position on roughly half the trials. This underscores an important caveat of the method: it may fail if the source model is incomplete.

**A two-dipole model.** The source model was modified to allow for two dipoles with equal amplitudes. The size of the two-dipole parameter space varies with the square of the number of dipole positions; for search to be feasible, this number was limited to 1000, chosen among positions of smallest cost function for the one-dipole model. Figure S8 C shows, for each location, the minimum of the 2-dipole cost function over all pairs with one dipole at that location (gray indicates positions outside the selected 1000). This pattern shows two clear minima at both source dipole locations. Figure S8 D shows a similar plot based on real MEG (as in Fig. S7 D). This simulation indicates that more complex sources can be handled with a more complete source model. However, this strategy is (a) costly for complex source spaces, (b) prone to "false positives" because a larger source space is easier to fit to the data. It is worth noting that this two-dipole model performs poorly if applied to a *one*-dipole source, or a *three*-dipole source (not shown).

This simulation underscores, again, the importance of the fit between the real source and the source model. It also highlights the tradeoff between flexibility (to accommodate all possible sources) and the cost and risk of spurious fits associated with a larger model space.

**Unconstrained dipole model.** The previous simulations were based on the constrained cortical surface model (dipoles oriented normal to that surface). Figure S9 A shows the cost function for an *unconstrained* cortical source model for the same source as in Fig. S7A. There is a clear minimum at the correct location, however for other source locations (about 16 %) the minimum falls at a different location. Figure S9 B (black dots) shows that most of those incorrect minima are within 5 mm of the correct position (average error 2.5 mm). The next-best estimate (red) and 3rd-6th best estimates are likewise quite close to the correct position. As this simulation shows, the flexibility of the unconstrained model leads to incorrect location estimates, however in this example the errors are quite small. The upside of the unconstrained model is that it can locate sources for which the constrained model is incorrect (incorrect orientation).

**How sensitive is the location estimate to imprecision of the forward model?** Figure S9 C shows a plot similar to Fig. S9 B, in the case where every coefficient of the gain matrix was jittered by adding a normally-distributed random value with an overall SNR of 10 (the variance of the added noise is 1 % of the RMS gain). Incorrect location estimates are more numerous (about 60 %) than in the absence of noise, however estimates remain clustered near the correct value (average error 7 mm). Figure S9 D shows the depth of the cost function minimum as a function of gain SNR for the constrained model. Imprecision in the forward model makes the estimates less accurate, but the degradation is not catastrophic.

**Multiple sources?** Figure S9 E shows the locations of ten sources. Each was excited with an independent Gaussian noise waveform, multiplied by the corresponding gain vector and added to other sources to obtain simulated MEG data. These were processed by PCA and columns 11-274 of the transform matrix were used as null filters, which were combined with the constrained model to obtain the cost function plotted in Fig. S9 F. Remarkably, the locations of all sources are recovered accurately, consistent with a similar result for the toy world (Fig. S5 B). This outcome is robust to correlations between the source time series, the only requirement being that they are not linearly dependent. If the source time series are *identical*, localization fails as in Fig. S8 B.

#### Real MEG

The method was tested with real MEG data recorded in response to repeated auditory stimulation (Supplementary Methods). The JD algorithm was applied to isolate three types of activity: time-locked to the stimulus, time-locked to the button press, or narrowband centered on 10 Hz (alpha) (Supplementary Methods). In each case, columns 6-274 of the JD transform matrix were used as null filters (columns 11-274 for alpha).

**Constrained cortical surface source model, auditory response.** Figure S10 A shows the cost function for null filters derived from the JD analysis time-locked to the auditory stimulus, based on the interval 0-200 ms post onset. For this analysis, sensors and model source locations were limited to the left hemisphere. Figure S10 B shows a similar analysis for the right hemisphere. The cost function takes large values (yellow) everywhere except within a zone located temporally on both sides.

The distribution of smaller values (blue or green) is roughly consistent with activation patterns obtained with other localization methods (dipole fitting, beamforming, and minimum norm estimate) as reported for the Brainstorm tutorial (not reproduced here). It must be noted, however, that those patterns that reflect *source activity*, whereas those of Fig. S10 reflect *goodness of fit* between the source model (dipole on the cortical surface, normal to that surface) and the real source. If the fit were perfect, the cost function would reach zero at some point in source space; the fact that it does not implies that *the real source does not fit the model*, i.e. it is not a single dipole located on the cortical surface and normal to that surface (according to the model). This mismatch could have multiple origins.

**Button press and alpha response.** Figure S10 C shows the cost function for a similar analysis time-locked to the button press, based on a 500 ms interval prior to the button press. Here, the location is frontal (consistent with motor cortex) and left-lateralized. However, the cost function dip is shallow, indicating a poor match the source model. Figure S10 D shows the cost function for a JD analysis designed to isolate alpha components. Here, the distribution of small values is barely visible because deep (it is easier to see on subsequent analyses).

**Two dipole model.** Figure S10 E-H shows similar results for a two-dipole version of the constrained cortical surface source model. The cost function is calculated only for 1000 model source locations (others are plotted in gray). Relatively large cost function values imply that the real source does *not* consist of two dipoles constrained to the cortical surface and normal to that surface.

**Unconstrained cortical surface source model**. Figure S10 I-L shows similar results for an unconstrained version of the cortical source model (dipole orientation is free). Again, the relatively large cost function values imply that the real source does *not* consist of a single dipole constrained to the cortical surface with arbitrary orientation.

**Volume source model.** Figure 4 of the main paper shows results similar to Fig. S10 for a volume source model (locations on a 2mm grid within the brain volume, arbitrary orientation). Whereas in Figs. S7-S10 only the foremost sources were visible, in Fig. 4 each pixel represents the minimum over a stack of voxels perpendicular to the image plane (“transparent brain”). With this difference in mind, patterns are overall consistent with the surface models.

The patterns in Fig. 11 A-G show considerable small-scale detail. For example, patterns reflecting auditory activity differ between left and right hemispheres, with an unexpected additional locus on the lower margin of the temporal lobe (also visible in Fig. 10 J). Alpha activity likewise shows a complex asymmetrical and multifocal pattern, mainly occipital but with hints of a more frontal distribution. Whether or not these details are veridical (representing the true distribution of cortical sources) is *unknown* at this point. These patterns must be interpreted with caution pending more complete validation.

**Comparison with other methods**. Figure S11 A shows patterns of activity over cortex at 110 ms post stimulus onset for six methods based on a 15002-dipole surface model. The first method assumes that dipoles are perpendicular to the cortical surface (constrained), the others assume arbitrary orientations (unconstrained). Figure S11 C shows similar patterns for five methods based on a model with dipoles distributed within the cortical volume on a grid with 2 mm pitch. Figure S11 B shows the cost function for the zero-set method. All plots are of a "transparent brain" type: each pixel in A and B represents the *maximum* of activity over a stack of voxels oriented perpendicular to the viewing plane, each pixel in B represents the *minimum* cost over a stack of voxels oriented perpendicular to the viewing plane. Plots in A and C show largest activity (yellow) in a temporal region (the measure and units differ between methods), consistent with the minimum of cost in panel C.

There are considerable differences in detail between methods, but in the absence of ground truth, one cannot conclude that any method is superior to others based on these plots. A major difference is that plots in A and C represent *activity* at a particular time (110 ms post stimulus onset), whereas B represents *cost* for a particular component chosen to maximize early evoked activity.

**Interictal spikes in EEG.** 29-channel EEG data from the Brainstorm tutorial on analysis of EEG in epilepsy were analyzed with JD time-locked to the interictal spikes (labeled within the data). Figure S11 H shows the time course of the first three JD components averaged over spikes (these components were further rotated so that the earliest activity is concentrated in the first component, and the temporal alignment of spikes was readjusted to maximize correlation over trials). Figure S10 I-K shows the cost function calculated from columns 2-29 of the JD matrix, using a volume model with a 2mm grid calculated with Brainstorm.

The cost function is minimum at a superficial location that appears centered on the left margin of the longitudinal fissure. This location is roughly consistent with locations determined by other methods (dipole fitting, beamforming, minimum norm estimate) as described in the Brainstorm tutorial (not reproduced here), albeit more precise. Whether this precision is real or illusory is *unknown* at this point. Once again, these patterns must be interpreted with caution pending more complete validation.

**Interictal spikes in SEEG**. Data from a patient implanted with 222 stereotactic EEG (SEEG) electrodes were obtained from a publicly-available dataset associated with a Brainstorm workshop (https://neuroimage.usc.edu/brainstorm/WorkshopParkCity2024). A head model with 126621 dipoles with arbitrary orientation distributed within the cortical volume at 2 mm intervals was prepared with Brainstorm. A 10 s recording including a labeled interictal spike was preprocessed (power line removal with ZapLine, dimensionality reduction with PCA, 2nd-order Butterworth high-pass filter with cutoff 200 Hz) and submitted to a JD analysis to emphasize power within the earliest portion of the spike (-100 to -20 ms relative to the marker). Columns 3-222 of the JD matrix were used as null filters. Figure S12 A-C shows the cost function in axial, sagittal and coronal views, together with electrode tracts. Figure S12 D-F shows an expanded view centered upon a minimum of the cost function.
The location of the minimum is within a zone well sampled by the electrode arrays, close to the electrodes most highly-correlated with the spike (as reflected by the diameters of the open symbols), but slightly offset from those electrodes. This offset suggests that ZSI refines the spatial information provided by the spatial pattern of spike amplitude over electrodes, however in the absence of ground truth this conclusion should be tempered.

**Summary of results**. Simulations show that the method can accurately localize sources within the parameter space of a realistic model, as long as they are consistent with that source model. Sources that do not initially fit can be handled by augmenting the source model, with two caveats. First, a larger source parameter space may be computationally expensive to search. Second, a larger model is more prone to false positives, or ambiguity.

On real data, the new method yielded estimates roughly consistent with estimates from other methods, and with expectations from prior knowledge. There are differences in detail (as there are between alternative methods), but it is not possible adjudicate between them at this stage for lack of ground truth.

An important issue is that, with real data, the method generally failed to find a parameter value for which the cost was zero, thus implying that the real source did *not* match the source model for any of its parameters. In principle, this could be overcome by augmenting the source model, in practice the search space is too large unless progress in understanding allows an exact model to be formulated. This being the case, it is tempting to interpret the pattern of (non-zero) cost over the source parameter space, or at least the position of its minimum, as informative (as in Fig. 5). Several results (with simulated or real data) seem to support this interpretation, but in the absence of theoretical basis caution is required.

### Supplementary figures

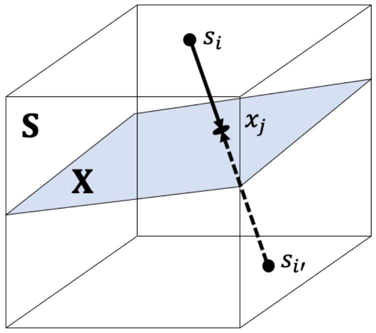


Fig. S1. Measurement of brain activity is a many-to-one transform, not invertible. Source activity (matrix $\mathbf{S}$) spans the vector space of their linear combinations, that includes observations (matrix $\mathbf{X}$) that span a subspace. Different sources may give rise to the same observation.

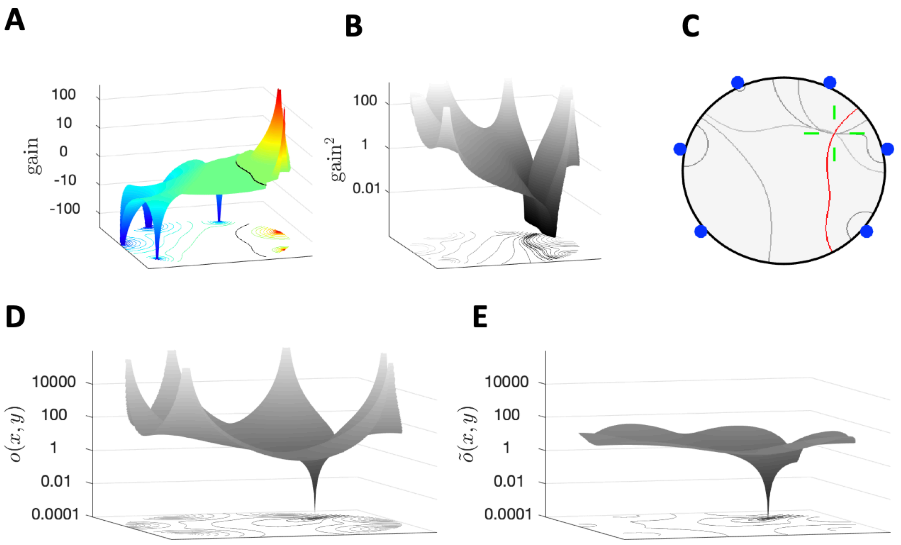


Fig. S2 A: Gain map of a spatial filter as a function of position within a 2D disc plotted in 3D and with a color scale. Negative values are plotted as -log10(-gain). The black line corresponds to the zero set of this filter. B: Squared gain of the same filter. The valley follows the zero set. C: Zero sets for five spatial filters that are all null filters with respect to a source situated at their intersection (green cross). D: Sum of squared gain maps for the five filters in C (Eq. S3). The minimum coincides with the location of the source. E: same, with normalization (Eq. S4).

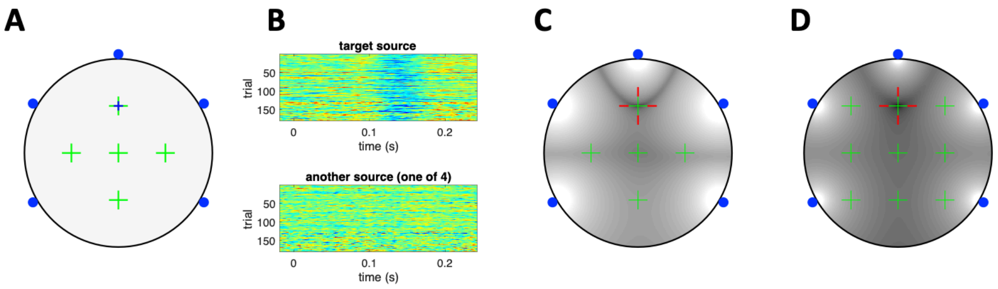


Fig. S3. Source localization using the JD algorithm. A: source positions are indicated as crosses (the blue cross is the target). B, top: waveform of the target source for individual trials plotted as a color scale. Bottom: same for a non-target source (one of 4). C: sum of squared gain maps of the null filters derived from columns 2-5 of the JD matrix. The position of the minimum of this pattern (red cross) correctly indicates the location of the target. D: same, for nine sources ($I > J$).

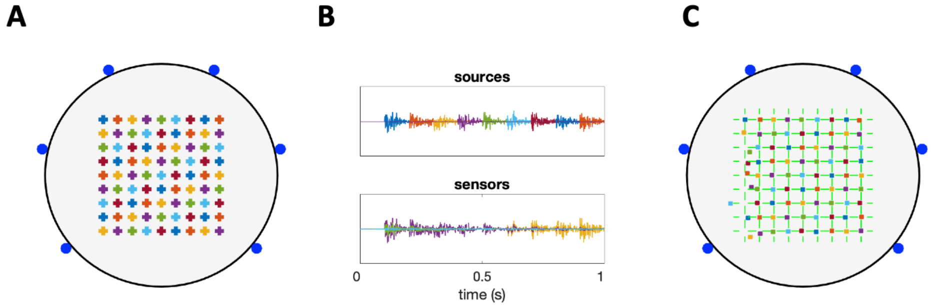


Fig. S4. Locating more sources than sensors. A: Source positions are indicated as crosses. B, top: source waveforms, colored as in A, bottom: sensor waveforms (arbitrary colors). C: estimated locations, colored as in A. The faint green crosses indicate the correct locations.

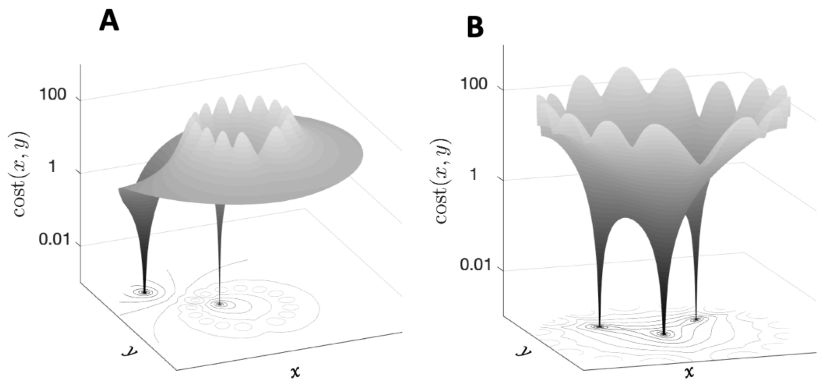

Fig. S5. A: Cost function for a single source and 15 sensors. For certain locations, a “ghost zero” appears. B: Cost function for 3 sources and 15 sensors (sum of squared gain maps of 12 null filters, normalized). Three zeros allow the three sources to be localized with permutation ambiguity.

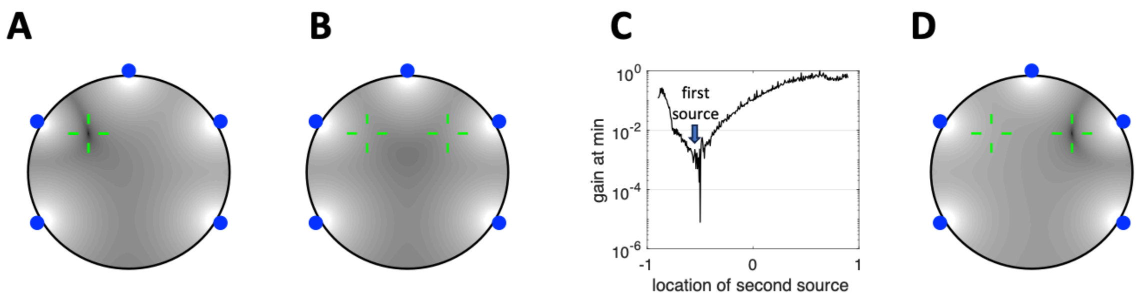


Fig. S6. A: Cost function values for a single-point source (green cross). B: Cost function values for a symmetric two-point source (green crosses). C: Value at the minimum of the cost function for a two-point source as a function of the second source location. D: Cost function for a two-point source model. Here, one point of the two-point model is fixed at the left cross, and the other point is free. The cost is minimal when the second point of the model source coincides with the right cross.

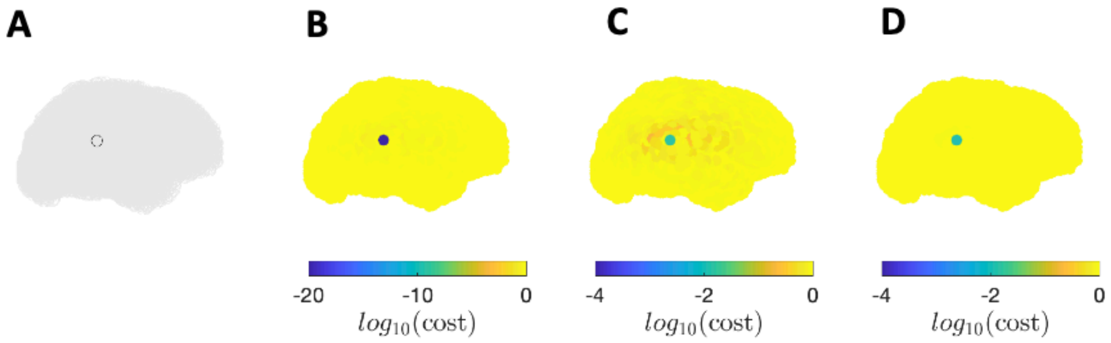


Fig. S7. Simulation of source localization with real MEG source and forward models. A: Location of a dipole source of Gaussian noise. B: Plot of the cost function based on null filters derived using PCA. C: Same, in the presence of channel-independent Gaussian noise at overall SNR=0.1 (-20 dB). D: Same, but target is an auditory-evoked MEG response (see text), background is phase=scrambled MEG at SNR=0.23 (≈-15dB), null filters are derived using JD.

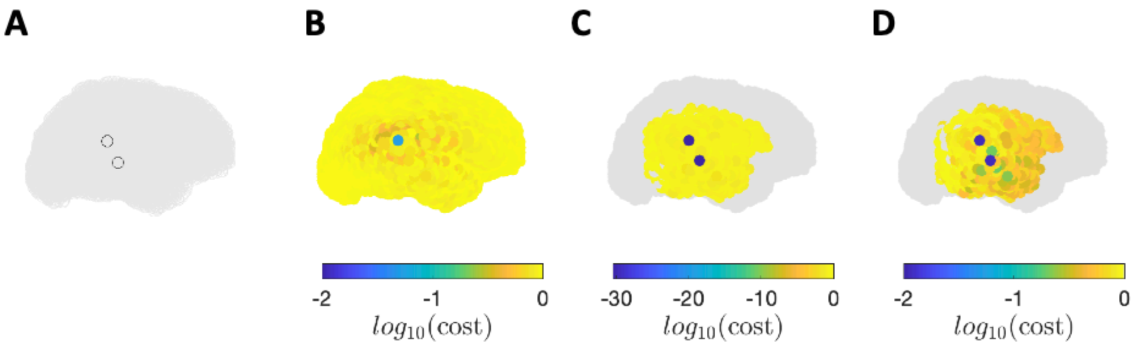


Fig. S8. A: Locations of a two-dipole source of Gaussian noise. B: Plot of the cost function over a one-dipole model space. C: Plot of the cost function over a two-dipole model space (see text). D: Same, but target is an auditory-evoked MEG response, as in Fig. S7.

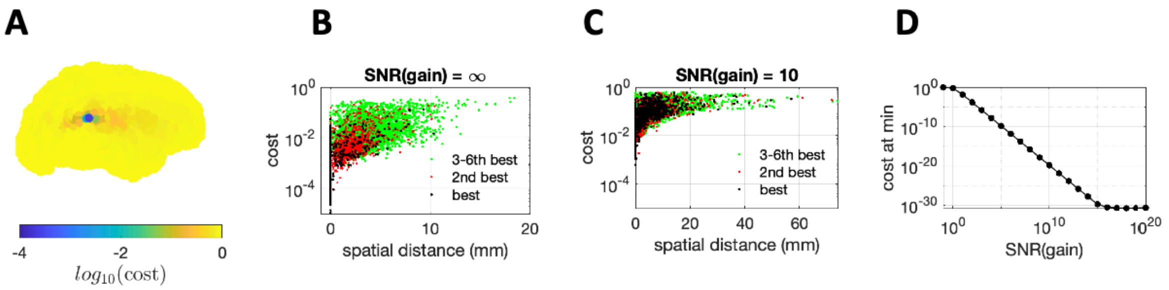


Fig. S9. A: Cost function for a dipole source fit by an unconstrained dipole model. B: Gain as a function of distance from the correct position of the best estimate (lowest cost, black), second best (red) and 3rd-6th best (green) for random sample of 1000 source locations. C: same as B in the case of a noisy gain matrix. D: depth of the overall min of the cost function as a function of the SNR of the gain matrix.

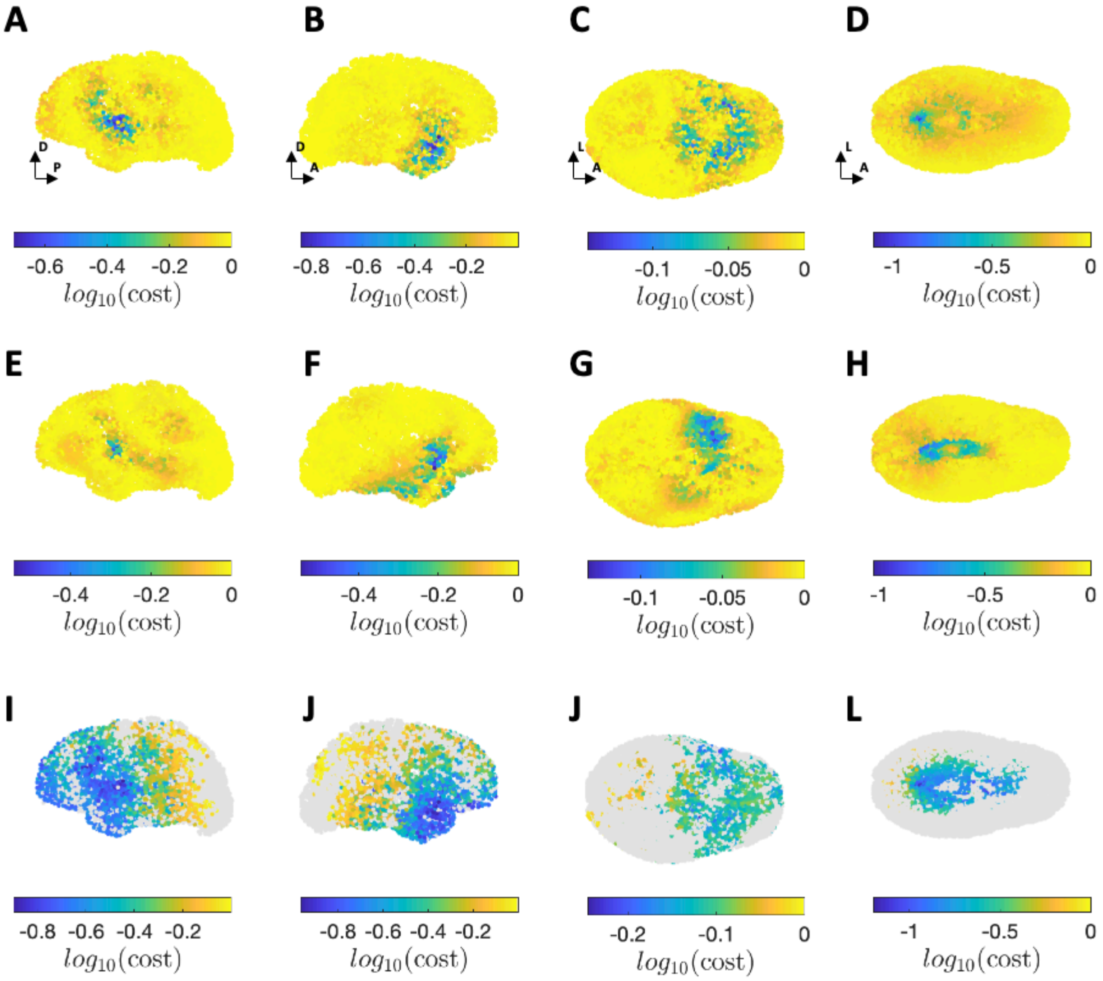


Fig. S10. Real MEG, alternative source models. A-D: cost function for constrained cortical surface source model. E-H: same, unconstrained. I-L: 2-dipole variant of constrained cortical source model. Compare with Fig. 4 in main paper. Deep sources are not visible in this representation.

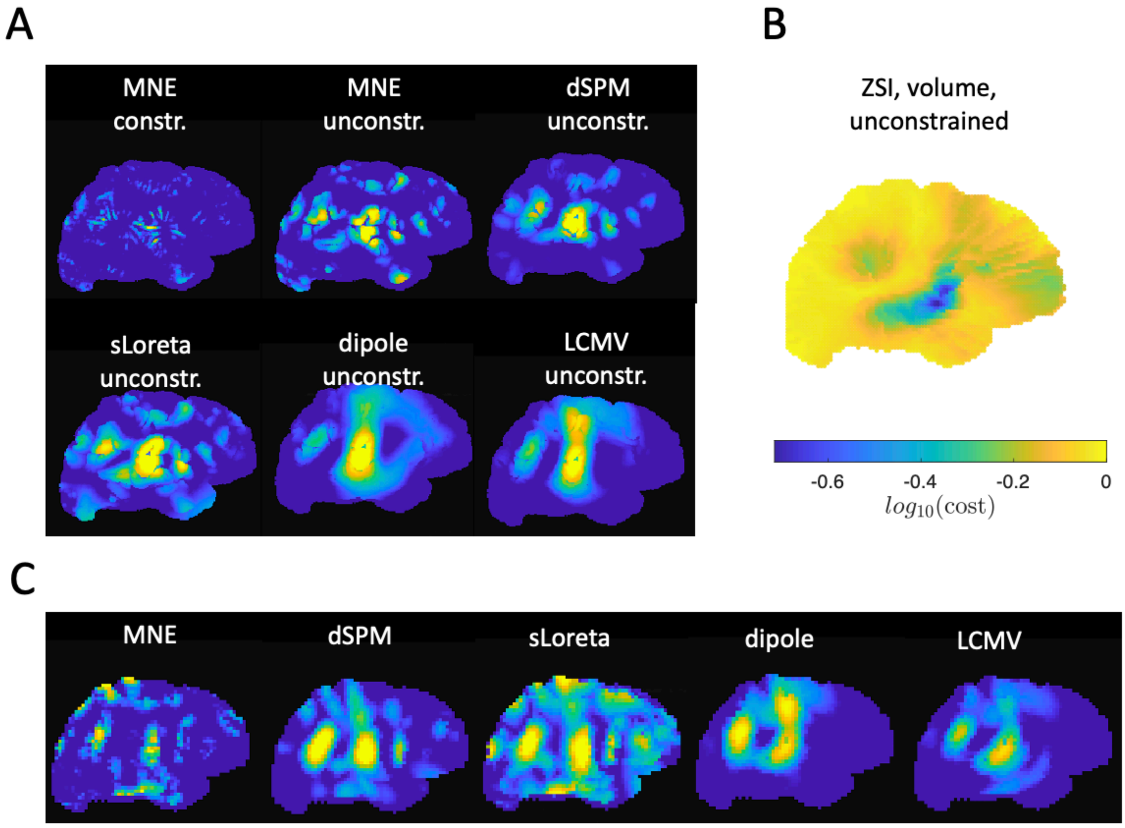


Fig. S11. Comparison with standard models, auditory stimulus-locked response. A: Activity patterns at 110 ms post-onset for six cortical surface models, implemented in Brainstorm. B: Cost function for the zero-set method. C: Activity patterns at 110 ms for five cortical volume models.

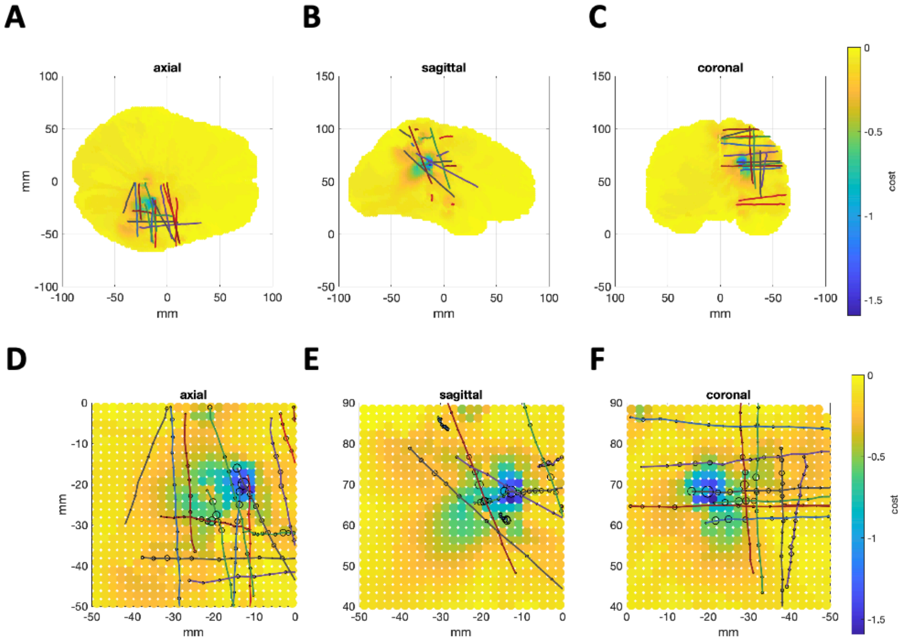

Fig. S12. Interictal spike in SEEG. A-C: cost function for null filters obtained from a JD analysis biased to enhance the early onset of an interictal spike (-100 to -20 ms relative to spike marker, data high-pass filtered with 200 Hz cutoff), in axial, sagittal and coronal views. Colored lines indicate electrode groups, markers indicate electrode positions. D-F: same, enlarged. The diameter of each open symbol is proportional to the correlation between that electrode and the selected JD component.
